# An ultrasensitive microfluidic approach reveals correlations between the physico-chemical and biological activity of experimental peptide antibiotics

**DOI:** 10.1101/2021.09.08.459503

**Authors:** Jehangir Cama, Kareem Al Nahas, Marcus Fletcher, Katharine Hammond, Maxim G. Ryadnov, Ulrich F. Keyser, Stefano Pagliara

## Abstract

Antimicrobial resistance challenges the ability of modern medicine to contain infections. Given the dire need for new antimicrobials, peptide antibiotics hold particular promise. These agents hit multiple targets in bacteria starting with their most exposed regions – their membranes. However, suitable assays to quantify the efficacy of peptide antibiotics at the membrane and cellular level have been lacking. Here, we employ two complementary microfluidic platforms to probe the structure-activity relationships of two experimental series of peptide antibiotics. We reveal strong correlations between each peptide’s physicochemical activity at the membrane level and biological activity at the cellular level by assaying the membranolytic activities of the antibiotics on hundreds of individual giant lipid vesicles, and quantifying phenotypic responses within clonal bacterial populations with single-cell resolution. Our strategy proved capable of detecting differential responses for peptides with single amino acid substitutions between them, and can accelerate the rational design and development of peptide antimicrobials.

## Introduction

Antimicrobial resistance is the “silent” pandemic that threatens to undermine modern medicine. Without intervention, it is predicted to cause 10 million deaths per year by 2050^1^, with losses to the global economy of up to $3.4 trillion p.a. within the next decade^2^. A combination of scientific and economic challenges^3–5^ has led to the drying up of the existing antimicrobial pipeline. Given the vital role that antimicrobials play in modern healthcare, it is crucial to develop and maintain new pipelines of effective antimicrobials, particularly with drugs that can circumvent currently prevalent resistance mechanisms.

Antimicrobial peptides (AMPs) have attracted significant interest as alternatives to traditional small molecule antibiotics^6–9^. In contrast to conventional antibiotics, there is no widespread resistance to these evolutionarily conserved molecules. AMPs are killing factors of innate immune systems of all life forms, which modulate host immune responses to clear infections^10^. Bacteria that are resistant to widely employed antibiotics have been shown to have a high frequency of collateral sensitivity to AMPs^11^. AMPs also show anti-biofilm activity, besides having anti-inflammatory and wound-healing properties^12^. In particular, cationic peptides have been designed to mimic the antimicrobial activity of α-helical host defence peptides that form an important part of the antimicrobial defences of multicellular organisms. These cationic peptides selectively target the anionic membranes of bacterial pathogens^13,14^. Encouragingly, a recent update from the CARB-X antibiotic accelerator reports that direct-acting peptides account for approximately 11% of their portfolio, showing renewed industry interest in these compounds to overcome multidrug resistant bacterial strains^15^.

However, the clinical translation of peptide antimicrobials has suffered from a lack of suitable assays to characterize their efficacy^16^. Standard bulk techniques such as Minimum Inhibitory Concentration (MIC) assays fail to uncover heterogeneities in drug activity within a population of genetically identical cells. Such cell-to-cell variability is a particular challenge when assessing the antimicrobial efficacy of peptides due to the inoculum effect^17^. Charged peptides can remain bound to cellular components after cell death and lysis, and hence affect the “free” peptide concentration^18^ observed by other cells in the vicinity, leading to heterogeneities in the response to the drug dose; this leads to some cells surviving the treatment in the absence of genetic resistance. Cationic peptides also adsorb to surfaces of glass and plastic, leading to a loss of free peptides from solutions^19^; this complicates experimental handling and may lead to inaccuracies in data interpretation in standard assays. A recent study by the Stella group confirmed a strong inoculum effect on the MICs of a range of antimicrobial peptides, which led the authors to question the utility of standard cellular screening assays for the field^20^. Other researchers have also identified a need for “new and standardized testing structures” to facilitate the translation of peptide antimicrobials from the bench to the clinic^16^. Given the fact that peptides multi-target, and in particular often attack bacterial membranes, characterization strategies should also include tests for membrane activity, in addition to cellular assays.

We recently reported the development of a novel set of ultrashort (≤ 11 amino acids) peptides, based on a minimal amphipathic helix with *bi*nary *en*coding (bien) by arginine and leucine, that showed antibacterial efficacy^21^. Notably, a single side-chain mutation in the hydrophobic face of the helix significantly changed the nature of peptide-lipid interactions leading to differential mechanistic and biological responses. An alanine at the mutation position (“bienA”) facilitated a deep insertion of the helix in lipid membranes, and the formation of circular pores in (anionic) supported lipid bilayers (SLBs). However, a lysine at the same position (“bienK”) led to a shallower insertion of the helix and the formation of fractal membrane ruptures in the SLBs (peptide sequences are provided in Table S1)^21^. To probe the connection between the biological (cellular) and physico-chemical (membranolytic) properties of these new peptides, here we study the interaction of the bienA and bienK peptides with hundreds of individual *Escherichia coli* cells using the microfluidic “mother-machine” device^22^, and in parallel investigate their interactions with model membranes in a bespoke assay on hundreds of giant unilamellar vesicles (GUVs)^23^. The use of these high-resolution platforms enabled us to reveal phenotypic variants that survived each of these new compounds and revealed important insights about their activity both at the membrane and cellular level, overcoming a number of the challenges associated with traditional screening assays.

## Results and Discussions

The “mother-machine” system enables the physical confinement of individual bacteria in well-controlled microenvironments, where they can be sequentially exposed to nutrients, drugs or viability indicators and the responses studied using time-lapse microscopy^24–26^. Briefly *E. coli* BW25113 cells were cultured overnight in Lysogeny Broth (LB) at 37°C to stationary phase and injected into the microfluidic device (see Experimental Section and ^24,25,27,28^). We waited for the cells to populate the “wells” of the chip (Figure 1), following which the trapped cells were challenged with a continuous flow of peptide for 3 h, with images taken hourly. After 3 h of treatment, the cells were provided with fresh LB to study survivors. Post overnight (O/N) LB treatment, dead staining was performed with propidium iodide (PI). Dead cells either lysed or took up PI and appeared bright under fluorescence illumination (Figure 1A). The survivors could be further categorised into those that were dividing (Figure 1B), and those that did not stain with the dye but did not divide either (Figure S1, Table S1) as previously reported^26–28^. As a caveat, we note that there is some debate in the literature about these non-dividing cells that do not stain with PI; some groups claim that they are simply dead cells^29^, while others have found them to be viable^26,28,30,31^. However, the cells that were dividing at the end of the experiment were unambiguously alive, and used for our quantitative analysis, as discussed below.

**Figure 1.**
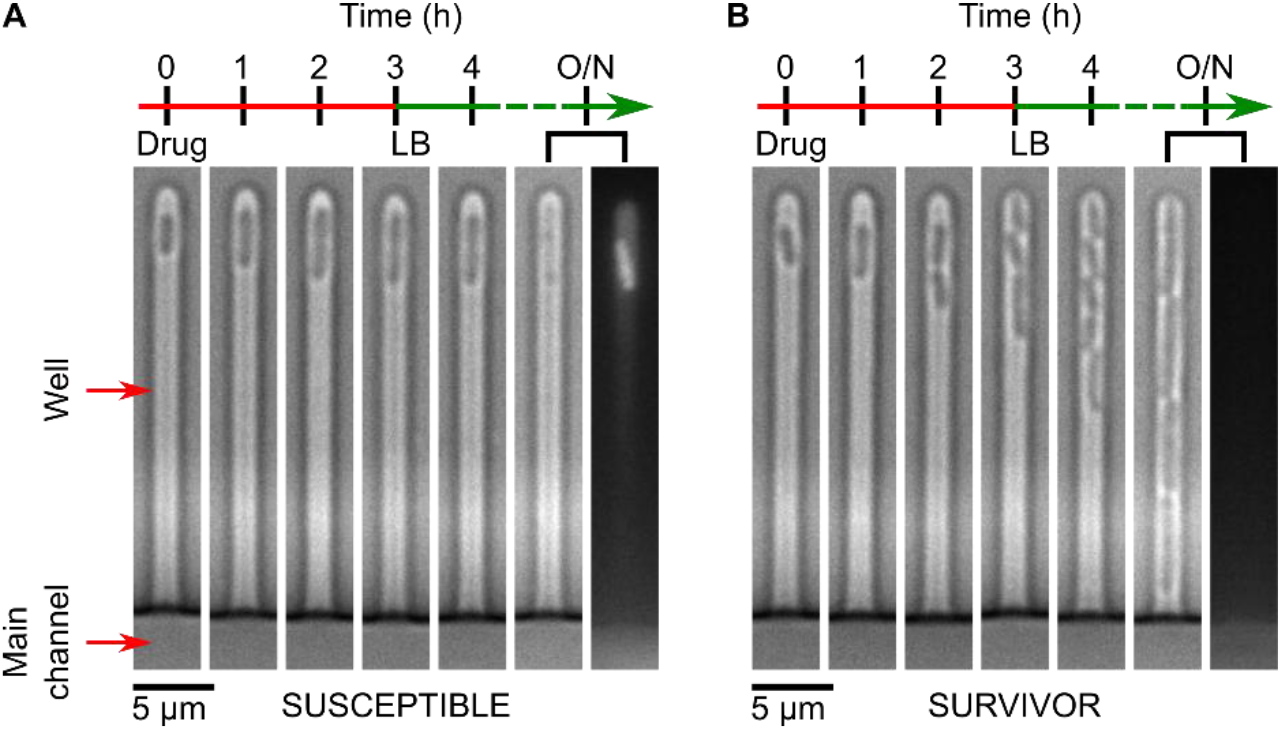
Probing the biological activity of experimental peptide antibiotics with single-cell resolution. In the images above, we observe examples of individual susceptible (A) and survivor (B) cells in response to peptide treatment in the mother-machine microfluidic device. The device consists of a “Main channel” for seeding the connected side channels or “wells” of the device with cells. Drugs, nutrients, and viability stains are then dosed to the trapped cells via the main channel. The two examples above are taken from the same experiment and show the contrasting responses of two clonal *E. coli* cells to the peptide bienK11 (10 μM). The cell shown in (A) was susceptible to the treatment, halting growth during the 3 h of drug dosage and disintegrating thereafter when the drug was replaced with fresh Lysogeny Broth (LB). In the final fluorescence image panel, taken after overnight (O/N) LB treatment, we see the debris of the cell stained with the dead stain propidium iodide (PI). In contrast, the cell shown in (B) resisted the treatment, growing and dividing even during peptide delivery, with no PI staining in the daughter cells at the end of the experiment.

We defined survival fraction as the fraction of wells hosting dividing cells after the 3 h peptide treatment and subsequent incubation in LB overnight (typically 17-18 h of LB incubation in total, see Experimental Section); as noted above, these dividing cells were unambiguously alive. This is equivalent to the survival fraction measured via colony forming unit (CFU) bulk assays and we therefore used this as a quantitative metric for the inhibitory efficacy of the peptides^26^. The bienA11 and bienA10 (hemolytic^21^) peptides were clearly the most potent in our assay, with a survival fraction of less than 5% (Figure 2A). In contrast, the bienA9 peptide was less effective with a survival fraction of 65 ± 11% (mean ± std. dev. over 2 biological repeats, 199 wells, 310 cells). Thus, reaching a critical peptide length leads to increased antibacterial activity with the bienA peptides. However, we did not find a similar dependence between antibiotic activity and peptide length for the bienK series, where we also observed a high frequency of phenotypic variants surviving peptide treatment. For bienK10 and bienK11, the survival fractions were 56 ± 26% (204 wells, 305 cells) and 31 ± 18% (215 wells, 311 cells) respectively, markedly different from the bienA10 and bienA11 results (Figure 2). Thus for these bienK peptides, while some individual cells were killed by the peptides, a large number of other clonal relatives survived the exact same conditions, similar to the example shown in Figure 1. Unlike traditional CFU or MIC assays, the peptides are dosed continuously for 3 h, and hence there should not be any significant differences around peptide exposure for the survivor versus susceptible cells and no inoculum effect. Our results reveal important phenotypic variants seemingly oblivious to the effects of peptide antibiotics that are able to grow and divide even during treatment, while their clonal relatives perish. We also observed similar trends when quantifying the cellular division rates in response to the peptides (Figure 2 C, D). The bienA11 and bienA10 peptides immediately and uniformly arrested cell division in all experiments, whereas bienA9 did not (Figure 2C), recapitulating the peptide length dependence observed for the survival fraction. Again, as with the survival fractions, this peptide length dependence was not observed for the cellular division rates with the bienK series (Figure 2D).

**Figure 2.**
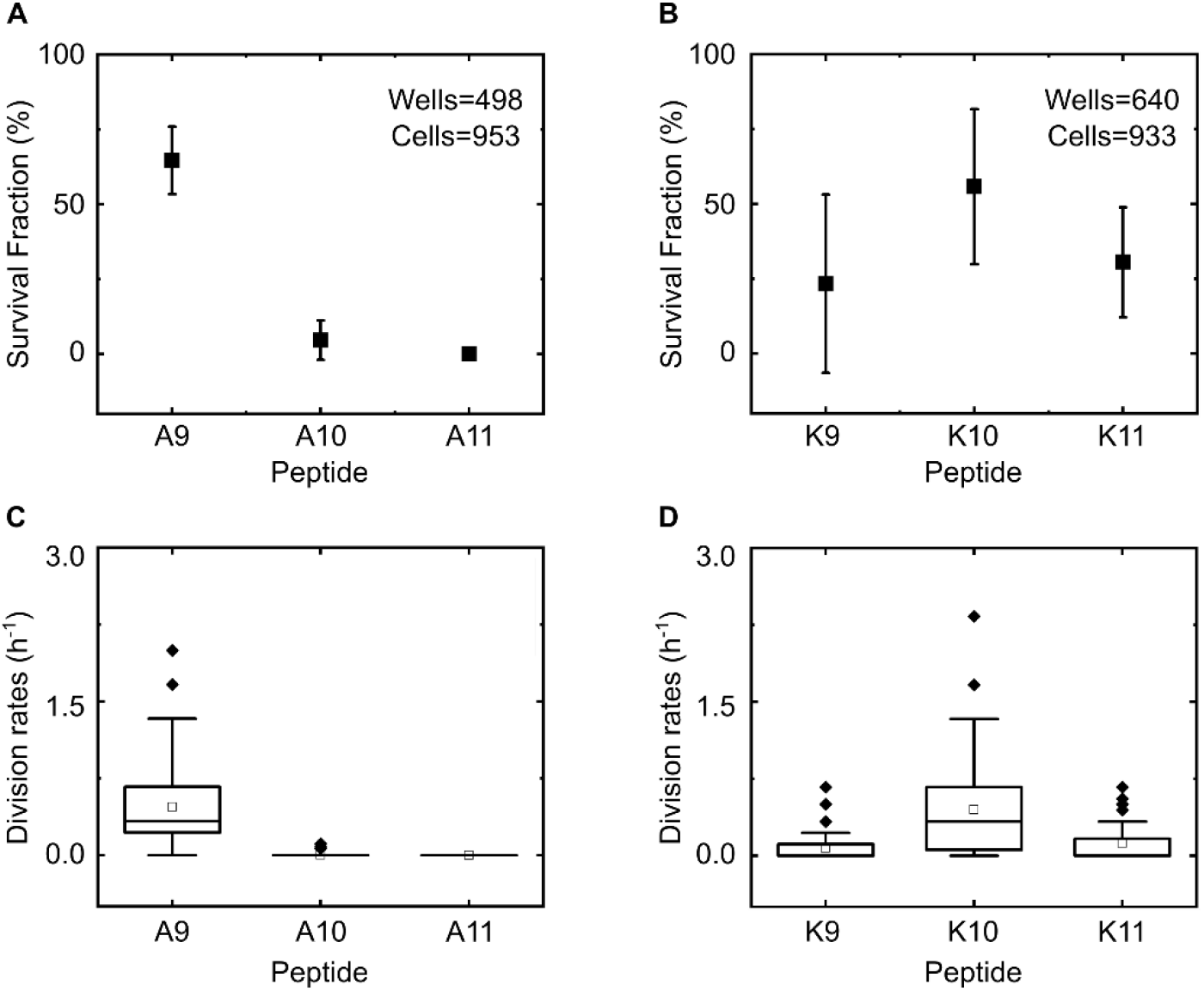
Single amino acid substitutions dramatically alter the efficacy of bien peptides at the single-cell level. From both the fraction of wells hosting dividing cells at the end of the experiment, (defined as the *survival fraction*, A, B) and the cell division rates during peptide treatment measured at the level of individual wells (normalized to the number of cells initially hosted by the well, C, D), we observed that the peptides bienA11 and bienA10 were the most potent of the set, with complete inhibition of division for bienA11 and a similar result for bienA10. In the bienA series, reaching a critical peptide length led to a marked increase in peptide activity, but this was not the case for the bienK series. BienA9 and bienK10 showed similar levels of activity, but were the least potent with over 50% of the wells showing cells that divided despite the treatment. BienK9 and bienK11 showed significant antimicrobial activity, but with considerable heterogeneity in the responses. Note, we typically only analysed wells that hosted between 1-3 cells each at the start of the experiment to avoid difficulties with tracking if the cells started growing, dividing, and exiting the channel during the drug treatment (this was not a concern for the bienA11 and bienA10 experiments since no/minimal cell division was observed in response to these peptides). Results are shown for two independent experiments (i.e., two independent culture isolates in independent technical repeats) for each treatment. We analysed at least 150 individual cells for each peptide experiment. All peptides were dosed at 10 μM concentrations. Plots in (A) and (B) report mean ± std. dev. For comparison, in our no-peptide control experiments, the survival fraction was 91 ± 11 % (162 cells tracked across 2 biological repeats).

Next, we sought an explanation for this phenotypic variability and investigated the mode of killing activity across the clonal population. When challenged with the bienA peptides, the cells showed rapid growth arrest; typically, no growth was seen after the 1 h observation time-point (Figure 3). In striking contrast, for cells treated with the bienK series, cells often elongated and septated until the point of division, when they appeared to stop growth and die (Figure 3A). This suggests a killing mechanism associated with the inhibition of processes involved in cell division. As noted in Figure 3B, cells treated with bienA10 and bienA11 showed a 20% increase in length on average before cell death, whereas with the bienK series cells doubled in length (on average) before death. This is possibly linked to the membrane activity of the bienK peptides, which were found in supported lipid bilayer (SLB) experiments to disrupt the outer leaflet of lipid bilayers^21^; these peptides may be exploiting weaknesses in the integrity of the membrane during the complex division process. However, there was significant variability in the response to each of the bienK peptides, and not all cells showed the same phenotype. For the cells analysed in Figure 3B, at the 3 h time point, the coefficient of variation (CV) in the lengths was 17%, 16% and 16% for the bienA9, bienA10 and bienA11 experiments respectively. For bienK9, bienK10 and bienK11, the CV values of the lengths were 44%, 47% and 48% respectively, showing the greater heterogeneity in lengths (at point of death) in response to each of the bienK peptides as compared to the bienA compounds. It is worth mentioning that the human antimicrobial peptide LL-37 has also been shown to attack and kill septating *E. coli* cells, where peptide binding was found to be preferential near the septum of the cells^32^. This suggests that there may be some similarities in the mechanisms of action of LL-37 and the bienK series of peptides.

**Figure 3.**
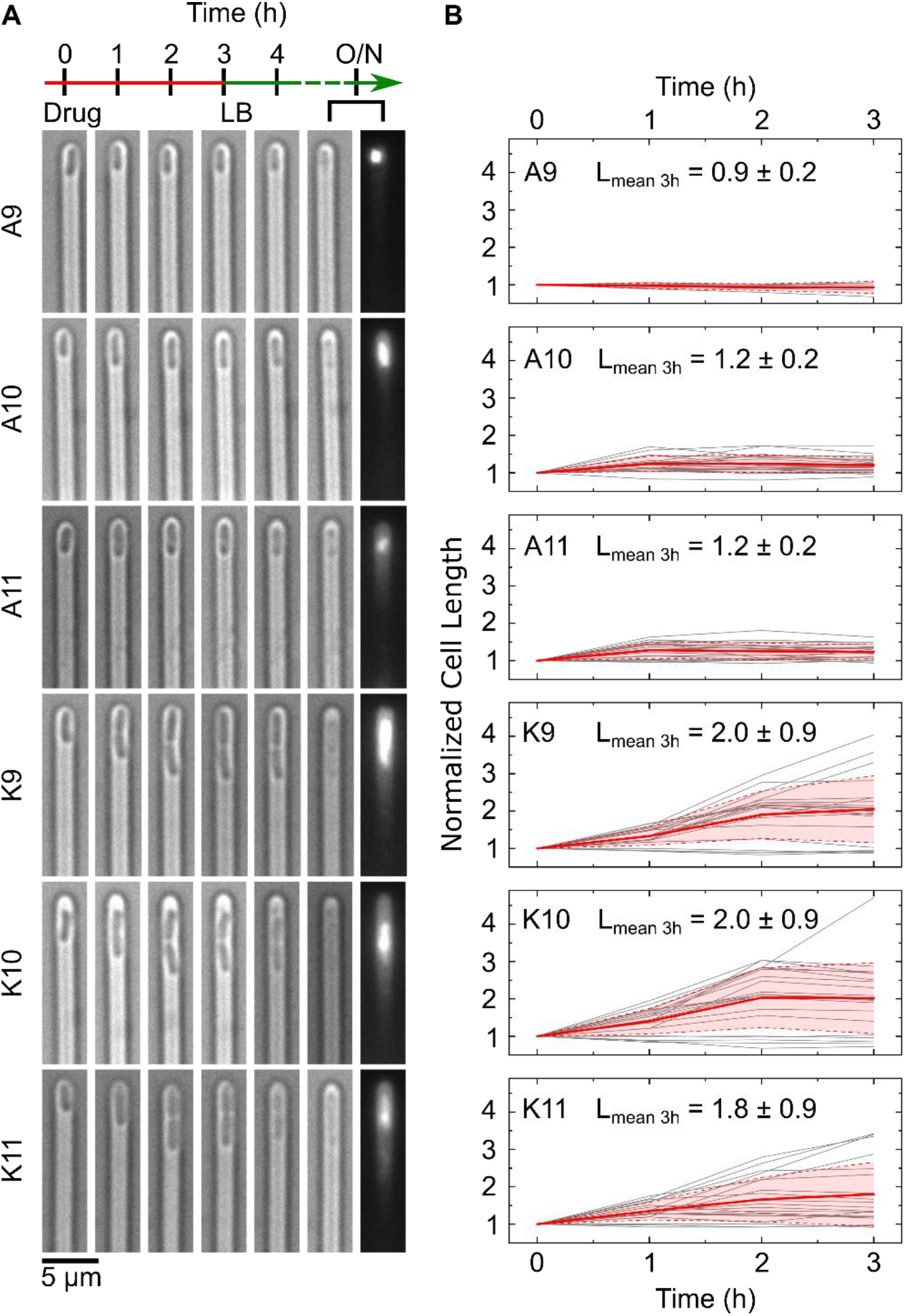
Comparing the single-cell killing phenotypes of our two experimental peptide series. The single-cell microscopy images in (A) depict differences in the killing phenotypes between the bienA and bienK peptides. The first image in each set (at *t* = 0) refers to the point at which the drug dosage starts, with the next images showing the cell at hourly intervals over 3 h of drug treatment (all peptides were dosed at 10 μM concentrations). The following image shows the cell after 1 h of LB incubation, followed by images showing the debris of the dead cell after overnight (O/N) incubation in LB (bright-field and corresponding PI stain fluorescence image). The images show that the bienA and bienK peptides kill *E. coli* cells in contrasting ways. The bienA series kill the cells rapidly, with the cells showing minimal elongation before growth arrest and death. This pattern is conserved across all the bienA peptides. This is also apparent from the corresponding graphs, shown in (B), tracking the lengths of individual cells (lengths are normalized to the initial cell length). Individual cell traces are provided in grey, with the mean (red) and standard deviation (red shaded area) depicted. The cellular length changes (before death) and distributions are relatively homogeneous for the bienA peptides. In stark contrast, the bienK peptides showed much greater heterogeneity in the lengths. Interestingly, for all three bienK peptides, we observed cells dying during septation, as depicted in the images in (A). However, as mentioned previously and as seen in the plots in (B), this was not the only phenotype. On average, with the bienK peptides, cells doubled in length before growth arrest and death, but there was significant spread in the lengths, as shown in the mean lengths (normalized) at the 3 h time-point reported inset in each plot (mean ± std. dev.). Note, for this analysis, we studied a subset of cells that were susceptible to the peptides, and were killed before undergoing any cell division. We chose a subset of 20 cells from 2 independent biological repeats for all the peptides, with the exception of bienA9 (5 cells, 1 experiment) which had far fewer cells showing the necessary characteristics (we excluded cells that lysed immediately after treatment, since their lengths could not be tracked over the entire course of drug treatment).

Given this variability in the cellular responses to treatment, we hypothesized that cell-to-cell differences at the membrane level could underlie the observed heterogeneity in the bacterial responses to these novel peptides. We therefore used anionic giant unilamellar vesicles (GUVs) as highly controlled model membranes to study the membranolytic activity of the bienA and bienK peptides^23^. The vesicles were produced in a microfluidic chip using octanol-assisted liposome assembly^33^ and made with a 3:1 mixture of 1,2-dioleoyl-sn-glycero-3-phosphocholine (DOPC) and 1,2-dioleoyl-sn-glycero-3-phospho-rac-(1-glycerol) sodium salt (DOPG) to mimic the anionic charge of bacterial membranes^23^. The GUVs were formed encapsulating a membrane impermeable dye, HPTS (8-hydroxypyrene-1,3,6-trisulfonic acid) and immobilized using an array of physical traps; the peptide solution was then flowed over them using a continuous flow system^23^. GUVs whose membranes are disrupted by the peptides lose their fluorescence and hence the system enables us to quantify the membranolytic efficacy of the peptides on hundreds to thousands of GUVs with single-vesicle resolution.

We performed experiments at both 5 and 10 μM peptide concentrations. Similarly to the single-cell studies, the single-GUV data showed striking differences between the bienA and bienK peptides (Figures 4, S2 and S3). In the bienA series, we observed a strong dependence of membranolytic efficacy on peptide length at the 5 μM concentration, with increasing peptide length increasing the membranolytic efficacy by up to 40-fold. In fact, vesicle survival rates (mean ± std. dev.) at 5 μM were 41 ± 3 % (bienA9, N=702), 12 ± 2 % (bienA10, N=808) and 1 ± 1 % (bienA11, N=813). At 10 μM, a similar trend was observed, with both bienA10 and bienA11 showing complete disruption of the entire vesicle population, while 19.9 ± 0.5% (N=559) of the vesicles survived the 10 μM bienA9 treatment. These results are in excellent agreement with the corresponding antibiotic activity observed in the single-cell experiments. In light of our previous studies with SLBs^21^, and the fact that the bienA peptides are hemolytic^21^, both our single-cell and single-GUV studies indicate a membranolytic mechanism of action for these peptides, with bienA11 being the most potent of the set.

**Figure 4.**
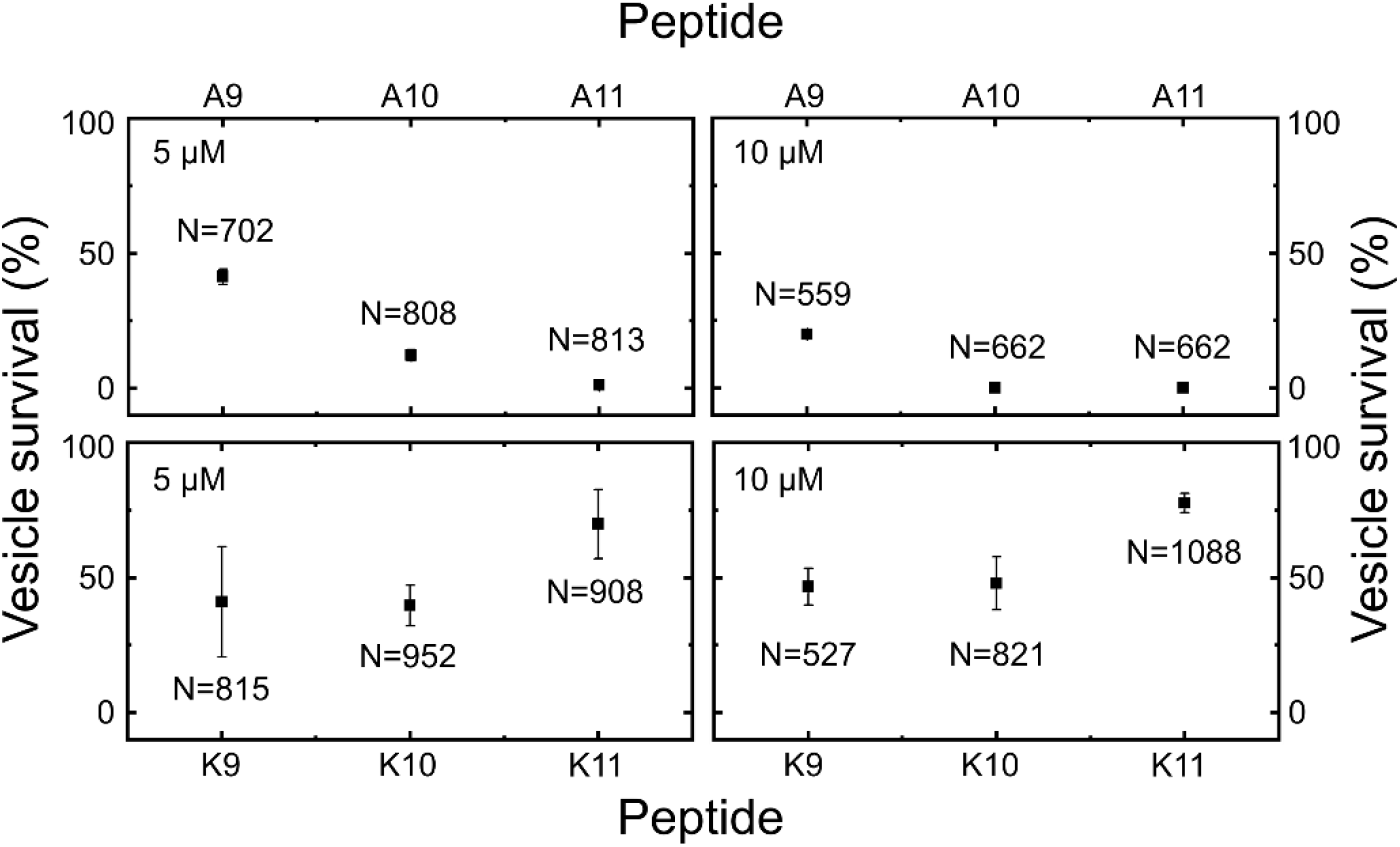
Vesicle viability levels after 7 hours of peptide administration. Data from two independent repeats per peptide, with total numbers of vesicles analysed mentioned inset (N). Data points represent mean survival percentages, and error bars report the standard deviations. Vesicles were continuously treated with the peptides at 5 μM or 10 μM concentrations in individual chambers each containing hundreds of physical traps. The vesicles in the control chambers were perfused with buffer. The majority of the population was stable under the flow of the control solution, with 86.8 ± 2.2 % of the vesicles surviving (mean ± std. dev., N = 633, 2 repeats). BienA11 was the most membranolytic peptide, followed by bienA10. At 5 μM, peptide activity increased with increasing length in the bienA series. At 10 μM, both bienA11 and bienA10 caused complete lysis of the vesicle populations. These trends are in agreement with the single-cell results reported in Figure 2. However, for the bienK series, none of the peptides led to complete lysis of the population of vesicles, even at 10 μM. Further, the activity of each of the bienK peptides was similar at both 5 and 10 μM concentrations, which indicates a different mode of action as compared to the bienA peptides. We hypothesise that the underlying antibiotic mechanisms for the bienK series involve membrane weakening, but potentially other intracellular mechanisms as well, based on the activity observed in cells. This requires further investigation. Please note, the single-GUV level data underlying this figure is provided in the SI (Figures S2 and S3).

In contrast, treatment with each of the bienK peptides led to considerable variability in vesicle survival within GUV populations, akin to the variability observed in the single-cell experiments. In the bienK9 and bienK10 experiments, at both 5 μM and 10 μM concentrations, over 40% of the GUVs survived the treatment with no dye leakage from the vesicle lumen (survival rates: bienK9: 41 ± 20 % for 5 μM, N=815; 47 ± 7 % for 10 μM, N=527; bienK10: 40 ± 7 % for 5 μM, N=952; 48 ± 10 % for 10 μM, N=821). For the bienK11 peptides, survival was even higher (survival rates: 70 ± 13 % for 5 μM, N=908; 78 ± 4 % for 10 μM, N=1088). This suggests that the bienK series is less membranolytic than the bienA peptides, which likely also explains the observation that the bienK peptides are not hemolytic^21^. This agrees with the mechanistic differences in membrane disruption by the two series observed in SLBs, where the bienA peptides formed transmembrane pores, while the bienK peptides caused fractal ruptures in the distal leaflet of the bilayer^21^. It is intriguing to speculate on the apparent membrane-to-membrane differences in the response to treatment by a bienK peptide, where we see a mix of GUV disruption and survival within a dosed population. Unlike cells, which display an inherent heterogeneity in gene, protein and lipid expression even within a clonal population, the GUVs being examined here all come from the same production batch within an experiment. These observations, coupled with the apparent affinity of bienK peptides for septating *E. coli* cells and the previous measurements on anionic SLBs^21^, suggest that the bienK peptides target bacterial membranes with a different mechanism to the bienA series, potentially exploiting weaknesses in the membranes of cells as they are undergoing division. Cell-to-cell heterogeneity in the lipopolysaccharides (LPS) of the *E. coli* outer membrane may also explain some of the heterogeneity observed in the single-cell experiments, since structural changes in LPS can confer resistance to cationic peptides^34,35^.

In addition to elucidating membrane-to-membrane differences in the membranolytic efficacy of a peptide within a vesicle population, we can also use our single-GUV resolution studies to reveal the *mode of action* of membrane-targeting antimicrobial peptides^36,37^. The mode of action is determined by studying the distribution of the time-points at which individual vesicles lose their fluorescence due to membrane disruption by the peptides, as previously described^36,37^. The heat-maps (SI Figures S2 and S3) demonstrate distinct time distributions of these single-GUV leakage events (LEs), which reveal the overall behaviour of the peptide-treated vesicle populations. In the cases of bienA11 and bienA10, the entire vesicle population exhibited LEs, meaning that all the individual GUVs’ membranes were disrupted (with 10 μM peptide). The LEs converged in a unimodal narrow distribution around a concentration-specific time-point. A similar trend was observed for bienA9, but in this case a subpopulation of vesicles survived at both treatment dosages. Based on the concentration dependent leakage behaviour, as well as the distributions of the timings of the LEs, we infer that the bienA series causes transmembrane disruptions that follow the two-state model described by Huang and others^37,38^. BienA11 and bienA10 thus subscribe to what is known as the unimodal “graded” mode of action while bienA9 follows the bimodal all-or-none mode^36^.

In contrast, in the bienK series, the LE data demonstrated similar membranolytic activity at both concentrations (Figures S2 and S3). The LEs followed a stochastic time distribution. Further, we observed high GUV survival rates in response to all three bienK sequences. Thus the mode of action for the bienK peptides is associated with stochastic LEs where a large fraction of GUVs survive the treatment. This behaviour has previously been observed in a 2-helix substructure of the peptide epidermicin^37^. In a manner similar to the bienK peptides, the epidermicin substructure also caused interfacial defects at the outer leaflet of a lipid bilayer rather than transmembrane disruption^21,39^, strengthening the correlation between this membrane thinning molecular mechanism and the stochastic LE mode of action^37^.

## Conclusions

Our single-cell and single-GUV level studies of experimental peptides reveal the importance of quantifying variability in the antimicrobial activity of individual peptides within a bacterial and GUV population. Characterizing this variability in response to treatment with a specific peptide – at both the membrane and cellular level - will be key for the further translational development of these candidate antibiotics. In particular, such membrane and cellular-level variability may translate into heterogeneous outcomes when such drugs are used to treat infections *in vivo*. Therefore, these peptide therapeutics will need to be carefully evaluated in pre-clinical models like those presented in this work to help design appropriate treatment strategies. More generally, our results add to the growing body of evidence that show some of the challenges associated with quantifying the activity of peptide-based therapeutics^16,20^. However, given the new strategy shown here and also complementary tools being developed elsewhere^20^, it is hoped that these difficulties can be quantified and addressed at an early stage of development, facilitating the efficient clinical translation of this promising category of therapeutics.

## Experimental Section

### Materials and Methods

#### Peptide synthesis

All peptides in the study were prepared as described elsewhere^21^. Briefly, the peptide sequences were assembled as peptide amides in a Liberty microwave peptide synthesizer (CEM Corp.) using conventional Fmoc/tBu solid-phase synthesis protocols on a Rink amide 4-methylbenzhydrylamine resin. After post-synthesis treatments, the peptides were purified using reversed-phase high performance liquid chromatography and identified by matrix assisted laser desorption ionization time of flight mass spectrometry.

#### Cell culture

The *Escherichia coli* (BW25113) cultures were prepared in 150 ml of Lysogeny Broth (LB; 10 g/L tryptone, 5 g/L yeast extract and 10 g/L NaCl, Melford) with shaking at 200 rpm and incubation at 37 °C overnight (drug dosage during experiments was typically performed around 19-20 h after starting the culture). Prior to seeding in the microfluidic chip, the OD of the culture was measured, the cells spun down at 3,220 g for 5 min and the supernatant filtered twice (0.22 μm, Millipore); this twice filtered supernatant is hereafter referred to as “spent LB”. The cell pellet was resuspended in spent LB at an approximate OD of 75; this concentrated culture was used when seeding the microfluidic chip with bacteria.

#### Microscopy for single-cell experiments

We used an Olympus IX73 epifluorescence microscope equipped with two Physik Instrumente piezo stages (M-545.USC and P-545.3C7) to control stage movement in XYZ. Images were acquired using an Olympus UPLSAPO 60×W objective (NA 1.2) and recorded on an Andor sCMOS (Zyla 4.2) camera with exposure times of 30 ms. We used the green LED of a pE-300 White LED light source (at 20% intensity) and a TRITC filter set for imaging the fluorescence of dead cells stained with propidium iodide (PI).

#### Peptide antibiotic preparation

Peptide stock solutions were always prepared fresh in milliQ water from the powders (stored at −80 °C) before each experiment. The absorbance of the stock solution was measured in a NanoDrop 2000 spectrometer at 214 nm, and used to calculate the stock concentration based on the corresponding extinction coefficient. We typically measured the absorbance of the stock solution at least thrice and used the average value to calculate the concentrations. For the cell experiments, the stock was diluted to 10 μM in a solution of 90% minimal media and 10% fresh LB (by volume) for use in the experiments^40^, with a total volume of 400 μl. The minimal media consisted of 1 × M9 salts, 2 mM MgSO_4_, 0.1 mM CaCl_2_ and 1 mg/L thiamine hydrochloride in milliQ water. For the control experiments, the dosage solution consisted of 30 μl of milliQ water (no peptide) in 370 μl of the minimal media-LB preparation. At each stage, the solutions were thoroughly vortexed to ensure complete mixing.

#### Microfluidics-microscopy assay for single-cell phenotyping

We recently published a detailed protocol of our microfluidics-microscopy assay, which is based upon the microfluidic “mother-machine” device pioneered by the Jun lab^25^; we have previously also used the device for quantifying survivor subpopulations in response to conventional small molecule antibiotics^26^. The device consists of a two-layer microfluidic chip with a main channel of width 100 μm and height 25 μm that is used for seeding the chip with bacteria, as well as nutrient, drug or dye dosing, along with thousands of smaller side channels, or “wells” (1.4 μm × 1.4 μm × 25 μm) which are used to physically confine the bacteria single-file for testing and long-term visualisation. The microfluidic chips are prepared from polydithmethylsiloxane (PDMS, Sylgard 184, Dowsil) by casting a 10:1 mixture of elastomer:curing agent (by weight) on an epoxy mold of the device (kindly provided by the Jun lab). This is cured for 2 h at 70 °C before being cut out, with inlet/outlet fluidic ports punched using a 0.75 mm biopsy punch (WellTech Rapid-Core). The chip is plasma bonded via a standard protocol (10 s plasma exposure, 30 W, Zepto plasma oven, Diener Electric, Germany) to a type I glass cover slip following which the channels are passivated with bovine serum albumin (BSA, 50 mg/ml in milliQ water) at 37 °C for at least 30 min. The chip is then seeded with a concentrated solution (OD595 approximately 75) of stationary phase *E. coli* via a syringe and tubing (Portex, microfluidic fine bore polythene tubing 1.09 mm × 0.38 mm). The chip is left for 5-10 min at 37 °C to allow the bacteria to enter the small side channels (wells) of the device, following which it is connected to the microfluidic pump system; we use a 4-channel Fluigent MFCS pressure pump with an associated Fluigent Flow Unit, which enables us to set a desired flow rate during the course of the experiments via a feedback system with the pressure pump. Experimental solutions are housed in the Fluigent Fluiwell-4C holder, which enables the exchange of solutions via a simple exchange of vials (1.5 ml, Micrewtube) without any disturbance to the microfluidic device itself.

After plugging the tubing (Fluigent) from the flow unit into the inlet of the chip and connecting the outlet to a waste reservoir, the flow unit is first calibrated and then used to flow spent LB for 8 min at 300 μl/h through the chip, to flush out the concentrated bacterial solution from the main channel of the device. Following this, the flow is stopped and the spent LB vial exchanged for a vial containing 400 μl of the peptide solution (10 μM). We chose 10 μM as the peptide concentration since this is above the reported MIC values (in the range of 3 - 6 μM) for these peptides in *E. coli* (ATCC 15597) cells^21^. To initially effect solution exchange, whenever the solution vial is changed, we flow the new solution at 300 μl/h for 8 min through the chip, and then reduce the flow rate to 100 μl/h. The chip is thus treated with the antibiotic for 3 h with bright-field images acquired at hourly intervals. Following 3 h of treatment, the antibiotic solution is exchanged with fresh LB (8 min at 300 μl/h followed by a 100 μl/h flow rate), with images again acquired for 3 h at hourly intervals. After 3 h, the flow is reduced to 50 μl/h and left overnight (typically for around 14-15 h). Finally, the cells are treated with the stain Propidium Iodide (PI, 1:1000 dilution of the stock in LB) for 8 min at 300 μl/h followed by 15 min at 100 μl/h, after which the cells are imaged in both bright-field and fluorescence mode (TRITC filter set, green LED) to characterise cell death and survival at the end of the experiment. Dead cells and debris stain with PI and show fluorescence, whereas cells that are seen intact in bright-field and remain dark under fluorescence illumination are classified as being alive. These include both survivors that divide and fill the channel as well as non-dividers that we have characterized previously^26–28^.

All image analyses were performed manually using FIJI. Note that, as shown in Figure 3, we often saw cells reaching the division point and then dying, which is why cells were only counted as having divided when it was clear that they were not still joined at the division site. This may in some cases lead to a slight underestimation of the division rate, but based on our experiments we noted that this was a more accurate representation of the results. This is also partly the reason we chose to analyse the images manually.

#### Microfluidics-microscopy assay for single-vesicle phenotyping

We have published detailed protocols for the design, fabrication and operation of the microfluidic device for immobilizing and testing thousands of individual GUVs, which can be found elsewhere^21,23^. Briefly, the platform facilitated three procedural steps integrated in a single device. First, the octanol-assisted liposome assembly^33^ technique was used to prepare GUVs. Next, the GUVs were trapped downstream in arrays of hydrodynamic posts. Finally, the immobilized GUVs were perfused with the desired dose of a peptide. All lipids described in the text were purchased from Avanti Polar Lipids. The GUVs were prepared in sucrose solution (200 mM) with glycerol (15% v/v) in PBS (pH 7.4). The vesicles were formed encapsulating the fluorescent membrane-impermeable dye 8-hydroxypyrene-1,3,6-trisulfonic acid from Thermo Fisher (HPTS, 50 μM). The microfluidic platform was operated by two positive pressure-driven pump modules (MFCS-4C, Fluigent), and a single neMESYS syringe pump module for fluid manipulation. The trapped GUVs were continuously dosed with buffer solution loaded with 5 μM HPTS (as a tracer) and the peptide of interest at 5 or 10 μM concentrations (the stock peptide solutions were prepared in milliQ water as described previously). Peptide arrival in the vesicle chamber, used to define the *t* = 0 time point, was monitored by tracking the HPTS tracer in the peptide perfusion solution. The data was later analysed using a custom Python code that detected the GUVs and collated their fluorescence intensity traces with peptide arrival in the microfluidic chambers.

## Acknowledgements

J.C. was supported by a Wellcome Trust Institutional Strategic Support Award to the University of Exeter (204909/Z/16/Z). This research was funded in whole, or in part, by the Wellcome Trust (Grant Number 204909/Z/16/Z). For the purpose of open access, the author has applied a CC BY public copyright licence to any Author Accepted Manuscript version arising from this submission. K.A.N. acknowledges support from a Cambridge-National Physical Laboratory (UK) studentship, the Winton Programme for the Physics of Sustainability, the Trinity-Henry Barlow Scholarship and the ERC. M.F. acknowledges support from an EPSRC iCASE studentship (reference 2148169, EP/R513180/1). U.F.K. acknowledges support from an ERC consolidator grant (Designerpores 647144). K.H. and M.G.R. acknowledge funding from the UK’s Department for Business, Energy and Industrial Strategy and Innovate UK (Grant Number 103358). S.P. was supported by an MRC Proximity to Discovery EXCITEME2 grant (MCPC17189), a BBSRC responsive mode grant (BB/V008021/1), a Royal Society research grant (RG180007), an award from the Gordon and Betty Moore Foundation Marine Microbiology Initiative (GBMF5514), and a Marie Skłodowska-Curie grant (H2020-MSCA-ITN-2015-675752).

## References

(1) O’Neill, J. Tackling Drug-Resistant Infections Globally: Final Report and Recommendations; 2016.

(2) World Bank. Drug-Resistant Infections: A Threat to Our Economic Future.; 2017.

(3) Wellcome. The Global Response to AMR: Momentum, Success, and Critical Gaps; 2020.

(4) Cama, J.; Henney, A. M.; Winterhalter, M. Breaching the Barrier: Quantifying Antibiotic Permeability across Gram-Negative Bacterial Membranes. J. Mol. Biol. 2019, 431 (18), 3531–3546.

(5) Cama, J.; Leszczynski, R.; Tang, P. K.; Khalid, A.; Lok, V.; Dowson, C. G.; Ebata, A. To Push or To Pull? In a Post-COVID World, Supporting and Incentivizing Antimicrobial Drug Development Must Become a Governmental Priority. ACS Infect. Dis. 2021, 7 (8), 2029–2042.

(6) Fjell, C. D.; Hiss, J. A.; Hancock, R. E. W.; Schneider, G. Designing Antimicrobial Peptides: Form Follows Function. Nat. Rev. Drug Discov. 2012, 11, 37–51.

(7) Lazzaro, B. P.; Zasloff, M.; Rolff, J. Antimicrobial Peptides: Application Informed by Evolution. Science 2020, 368, eaau5480.

(8) Koehbach, J.; Craik, D. J. The Vast Structural Diversity of Antimicrobial Peptides. Trends Pharmacol. Sci. 2019, 40 (7), 517–528.

(9) Grassi, L.; Maisetta, G.; Esin, S.; Batoni, G. Combination Strategies to Enhance the Efficacy of Antimicrobial Peptides against Bacterial Biofilms. Front. Microbiol. 2017, 8, 2409.

(10) Magana, M.; Pushpanathan, M.; Santos, A. L.; Leanse, L.; Fernandez, M.; Ioannidis, A.; Giulianotti, M. A.; Apidianakis, Y.; Bradfute, S.; Ferguson, A. L.; Cherkasov, A.; Seleem, M. N.; Pinilla, C.; de la Fuente-Nunez, C.; Lazaridis, T.; Dai, T.; Houghten, R. A.; Hancock, R. E. W.; Tegos, G. P. The Value of Antimicrobial Peptides in the Age of Resistance. Lancet Infect. Dis. 2020, 20, e216–e230.

(11) Lázár, V.; Martins, A.; Spohn, R.; Daruka, L.; Grézal, G.; Fekete, G.; Számel, M.; Jangir, P. K.; Kintses, B.; Csörgő, B.; Nyerges, Á.; Györkei, Á.; Kincses, A.; Dér, A.; Walter, F. R.; Deli, M. A.; Urbán, E.; Hegedűs, Z.; Olajos, G.; Méhi, O.; Bálint, B.; Nagy, I.; Martinek, T. A.; Papp, B.; Pál, C. Antibiotic-Resistant Bacteria Show Widespread Collateral Sensitivity to Antimicrobial Peptides. Nat. Microbiol. 2018, 3, 718–731.

(12) Haney, E. F.; Straus, S. K.; Hancock, R. E. W. Reassessing the Host Defense Peptide Landscape. Front. Chem. 2019, 7, 43.

(13) Sato, H.; Feix, J. B. Peptide–Membrane Interactions and Mechanisms of Membrane Destruction by Amphipathic α-Helical Antimicrobial Peptides. Biochim. Biophys. Acta - Biomembr. 2006, 1758, 1245–1256.

(14) Shai, Y. Mode of Action of Membrane Active Antimicrobial Peptides. Biopolym. Peptide Sci. 2002, 66, 236–248.

(15) Duffy, E. M.; Buurman, E. T.; Chiang, S. L.; Cohen, N. R.; Uria-Nickelsen, M.; Alm, R. A. The CARB ‑ X Portfolio of Nontraditional Antibacterial Products. ACS Infect. Dis. 2021, 7 (8), 2043–2049.

(16) Meurer, M.; O’Neil, D. A.; Lovie, E.; Simpson, L.; Torres, M. D. T.; de la Fuente-nunez, Cesar Angeles-boza, A. M.; Kleinsorgen, C.; Mercer, D. K.; von KocKritz-Blickwede, M. Antimicrobial Susceptibility Testing of Antimicrobial Peptides Requires New and Standardized Testing Structures. ACS Infect. Dis. 2021, 7 (8), 2205–2208.

(17) Jepson, A. K.; Schwarz-Linek, J.; Ryan, L.; Ryadnov, M. G.; Poon, W. C. K. What Is the ‘Minimum Inhibitory Concentration’ (MIC) of Pexiganan Acting on Escherichia Coli?—A Cautionary Case Study. In Leake M. (eds) Biophysics of Infection. Advances in Experimental Medicine and Biology, vol 915. Springer, Cham; 2016.

(18) Snoussi, M.; Talledo, J. P.; Del Rosario, N.-A.; Mohammadi, S.; Ha, B.-Y.; Košmrlj, A.; Taheri-Araghi, S. Heterogeneous Absorption of Antimicrobial Peptide LL37 in Escherichia Coli Cells Enhances Population Survivability. Elife 2018, 7, e38174.

(19) Kristensen, K.; Henriksen, J. R.; Andresen, T. L. Adsorption of Cationic Peptides to Solid Surfaces of Glass and Plastic. PLoS One 2015, 10 (5), e0122419.

(20) Loffredo, M. R.; Savini, F.; Bobone, S.; Casciaro, B.; Franzyk, H.; Mangoni, M. L.; Stella, L. Inoculum Effect of Antimicrobial Peptides. Proc Natl Acad Sci U S A 2021, 118 (21), e2014364118.

(21) Hammond, K.; Cipcigan, F.; Nahas, K. Al; Losasso, V.; Lewis, H.; Cama, J.; Martelli, F.; Simcock, P. W.; Fletcher, M.; Ravi, J.; Stansfeld, P. J.; Pagliara, S.; Hoogenboom, B. W.; Keyser, U. F.; Sansom, M. S. P.; Crain, J.; Ryadnov, M. G. Switching Cytolytic Nanopores into Antimicrobial Fractal Ruptures by a Single Side Chain Mutation. ACS Nano 2021, 15 (6), 9679–9689.

(22) Wang, P.; Robert, L.; Pelletier, J.; Dang, W. L.; Taddei, F.; Wright, A.; Jun, S. Robust Growth of Escherichia Coli. Curr. Biol. 2010, 20 (12), 1099–1103.

(23) Al Nahas, K.; Cama, J.; Schaich, M.; Hammond, K.; Deshpande, S.; Dekker, C.; Ryadnov, M. G.; Keyser, U. F. A Microfluidic Platform for the Characterisation of Membrane Active Antimicrobials. Lab Chip 2019, 19, 837–844.

(24) Cama, J.; Voliotis, M.; Metz, J.; Smith, A.; Iannucci, J.; Keyser, U. F.; Tsaneva-Atanasova, K.; Pagliara, S. Single-Cell Microfluidics Facilitates the Rapid Quantification of Antibiotic Accumulation in Gram-Negative Bacteria. Lab Chip 2020, 20, 2765–2775.

(25) Cama, J.; Pagliara, S. Microfluidic Single-Cell Phenotyping of the Activity of Peptide-Based Antimicrobials. In Polypeptide Materials: Methods and Protocols, Methods in Molecular Biology; Ryadnov, M. G., Ed.; 2021; Vol. 2208, pp 237–253.

(26) Bamford, R. A.; Smith, A.; Metz, J.; Glover, G.; Titball, R. W.; Pagliara, S. Investigating the Physiology of Viable but Non-Culturable Bacteria by Microfluidics and Time-Lapse Microscopy. BMC Biol. 2017, 15, 121.

(27) Goode, O.; Smith, A.; Łapińska, U.; Bamford, R.; Kahveci, Z.; Glover, G.; Attrill, E.; Carr, A.; Metz, J.; Pagliara, S. Heterologous Protein Expression Favors the Formation of Protein Aggregates in Persister and Viable but Nonculturable Bacteria. ACS Infect. Dis. 2021, 7, 1848–1858.

(28) Goode, O.; Smith, A.; Zarkan, A.; Cama, J.; Invergo, B. M.; Belgami, D.; Caño-Muñiz, S.; Metz, J.; O’Neill, P.; Jeffries, A.; Norville, I. H.; David, J.; Summers, D.; Pagliara, S. Persister Escherichia Coli Cells Have a Lower Intracellular pH than Susceptible Cells but Maintain Their pH in Response to Antibiotic Treatment. mBio 2021, 12 (4), e00909–21.

(29) Song, S.; Wood, T. K. ‘Viable but Non-Culturable Cells’ Are Dead. Environ. Microbiol. 2021, 23 (5), 2335–2338.

(30) Ayrapetyan, M.; Williams, T. C.; Oliver, J. D. Bridging the Gap between Viable but Non-Culturable and Antibiotic Persistent Bacteria. Trends Microbiol. 2015, 23 (1), 7–13.

(31) Henry, T. C.; Brynildsen, M. P. Development of Persister-FACSeq: A Method to Massively Parallelize Quantification of Persister Physiology and Its Heterogeneity. Sci. Rep. 2016, 6, 25100.

(32) Sochacki, K. A.; Barns, K. J.; Bucki, R.; Weisshaar, J. C. Real-Time Attack on Single Escherichia Coli Cells by the Human Antimicrobial Peptide LL-37. Proc Natl Acad Sci U S A 2011, 108 (16), E77–E81.

(33) Deshpande, S.; Caspi, Y.; Meijering, A. E. C.; Dekker, C. Octanol-Assisted Liposome Assembly on Chip. Nat. Commun. 2016, 7, 10447.

(34) Klein, G.; Lindner, B.; Brade, H.; Raina, S. Molecular Basis of Lipopolysaccharide Heterogeneity in Escherichia Coli. J. Biol. Chem. 2011, 286 (50), 42787–42807.

(35) Raetz, C. R. H.; Reynolds, C. M.; Trent, M. S.; Bishop, R. E. Lipid A Modification Systems in Gram-Negative Bacteria. Annu. Rev. Biochem. 2007, 76, 295–329.

(36) Nahas, K. Al; Keyser, U. F. Standardizing Characterization of Membrane Active Peptides with Microfluidics. Biomicrofluidics 2021, 15, 041301.

(37) Al Nahas, K.; Fletcher, M.; Hammond, K.; Nehls, C.; Cama, J.; Ryadnov, M. G.; Keyser, U. F. Measuring Thousands of Single Vesicle Leakage Events Reveals the Mode of Action of Antimicrobial Peptides. bioRxiv 2021, 455434.

(38) Huang, H. W. Action of Antimicrobial Peptides : Two-State Model. Biochemistry 2000, 39 (29), 8347–8352.

(39) Hammond, K.; Lewis, H.; Halliwell, S.; Desriac, F.; Nardone, B.; Ravi, J.; Hoogenboom, B. W.; Upton, M.; Derrick, J. P.; Ryadnov, M. G. Flowering Poration — A Synergistic Multi-Mode Antibacterial Mechanism by a Bacteriocin Fold. iScience 2020, 23, 101423.

(40) Pu, Y.; Zhao, Z.; Li, Y.; Zou, J.; Ma, Q.; Zhao, Y.; Ke, Y.; Zhu, Y.; Chen, H.; Baker, M. B.; Ge, H.; Sun, Y.; Xie, X. S.; Bai, F. Enhanced Efflux Activity Facilitates Drug Tolerance in Dormant Bacterial Cells. Mol. Cell 2016, 62, 284–294.

