## Supplementary Information for "An ultrasensitive microfluidic approach reveals correlations between the physico-chemical and biological activity of experimental peptide antibiotics"

Supplementary information includes:

- Table S1
- Figures S1-S3
- Supplementary references

| **Treatment (10 μM)** | **Peptide sequence (mutation position highlighted in red)** | **Initial no. of cells (*t*=0)** | **No. of non-dividing survivors at the end of the experiments** |
|---|---|---|---|
| Bien A9 | RLLRL**A**LRL | 310 | 11 |
| Bien A10 | RLLRL**A**LRLL | 320 | 6 |
| Bien A11 | RLLRL**A**LRLLR | 323 | 24 |
| Bien K9 | RLLRL**K**LRL | 317 | 13 |
| Bien K10 | RLLRL**K**LRLL | 305 | 7 |
| Bien K11 | RLLRL**K**LRLLR | 311 | 1 |

**Table S1. The peptides under investigation, with their corresponding amino acid sequences.** In response to our treatment (10 μM peptide), in addition to the results reported in the main manuscript, we observed a small number of *E. coli* cells that did not divide, but also did not stain with the dead-stain propidium iodide (PI) at the end of the experiments (Figure S1B). These cells are potentially survivors, akin to phenotypes we have reported previously[1–3].

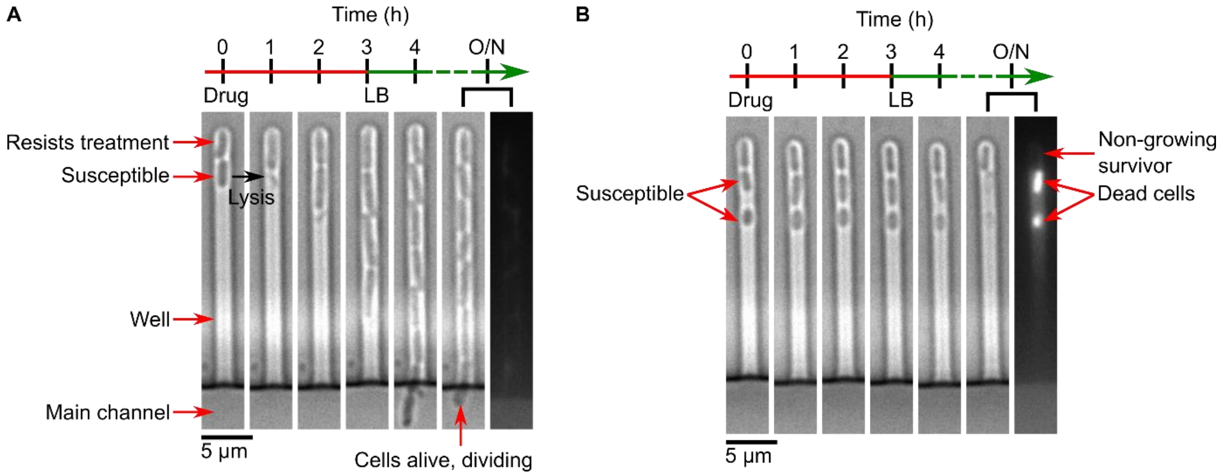

**Figure S1. Representative images showing the different cellular phenotypes observed in response to peptide treatment.** In (A), the images depict two *E. coli* cells trapped in a microfluidic well (at $t = 0$), subjected to 10 μM of the bienA9 peptide for 3 h, followed by fresh nutrient delivery (LB). The panel shows bright-field microscopy images taken at hourly intervals during peptide treatment, followed by bright-field images after 1 h and overnight (O/N) growth in fresh LB. The last panel shows the cells (fluorescence imaging) after treatment with the dead stain propidium iodide (PI). The initial two (clonal) cells are from the same culture and exposed to identical treatments. As can be seen, the cell at the bottom lysed upon exposure to the peptide. In stark contrast, its neighbour resisted the peptide, growing and dividing through the treatment. The daughter cells continued dividing thereafter in fresh LB media (cells post overnight growth were alive and did not stain with PI). In (B), we track the response of 3 individual, clonal *E. coli* cells to the bienA10 peptide (10 μM). None of the cells divided either during treatment or after fresh LB media was flushed through the device. However, only 2 of the cells died and stained with PI. These cells also disintegrated after the overnight LB treatment. In contrast, the topmost cell in the images did not disintegrate, nor did it stain with PI. Yet it did not divide, akin to the so-called “viable but non culturable” (VBNC) phenotype that we have characterized previously[1–3]. We label such cells “non-dividing survivors” for the purposes of this paper.

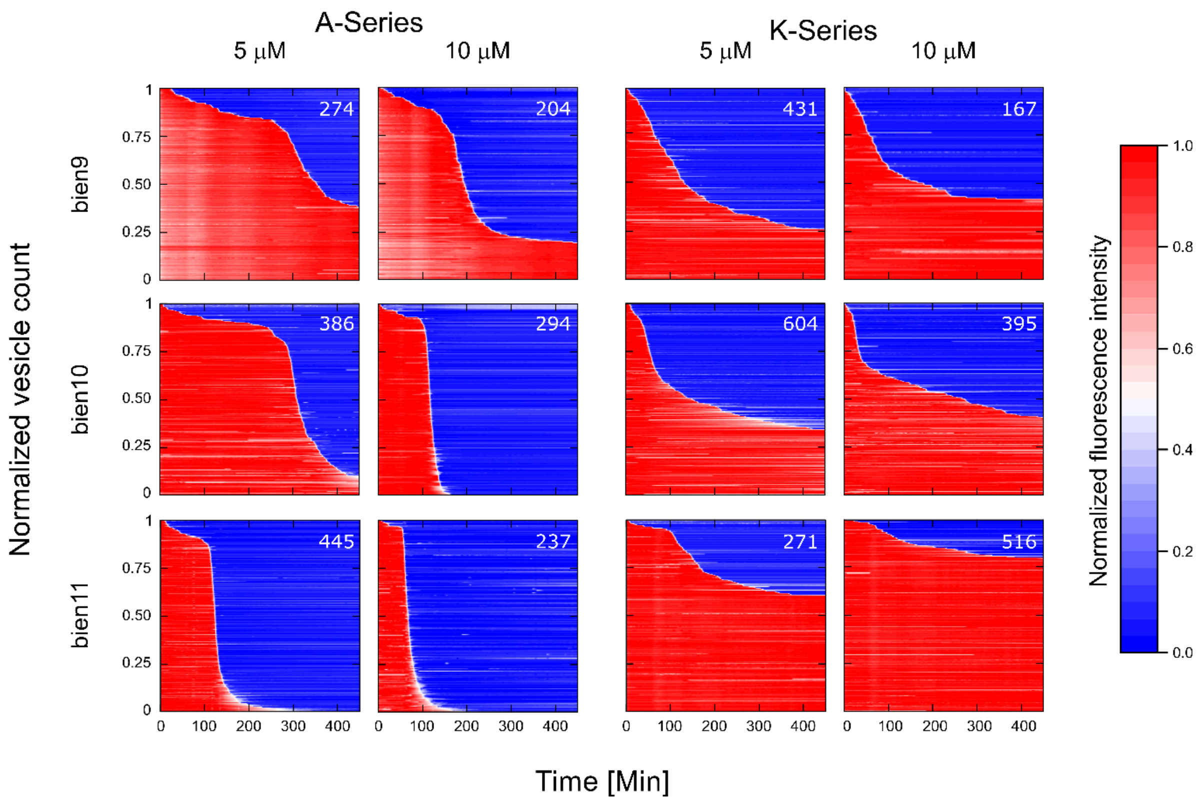

**Figure S2. A summary of the membranolytic activity of the bien peptides on populations of bacterial membrane-mimicking lipid vesicles. Dataset 1 of 2.** Trapped vesicles were continuously treated with the peptides at 5 µM and 10 µM concentrations and their morphology was observed overnight. Each horizontal line depicts the locally normalized intensity of the fluorescent dye HPTS encapsulated in a single trapped vesicle (with global background subtraction) over time. The vesicle's membrane is considered intact at high fluorescence intensity (red) and compromised at low fluorescent signal (blue). The intensity traces were ordered by the critical viability time point, which is defined as the point when the fluorescence intensity of a vesicle decreases below 50% of its initial intensity. The total number of analysed vesicles is reported in white in the top right corner of every plot. The results show that the bienA series of peptides is membranolytic, with potency increasing as one progresses from bienA9 to bienA11 and with an increase in the respective drug concentrations. However, the bienK series is not obviously membranolytic – there appears to be a weakening of the membranes in relation to controls, but there is no obvious concentration dependence, and the trend is reversed with respect to the bienA series, with bienK11 being the least potent.

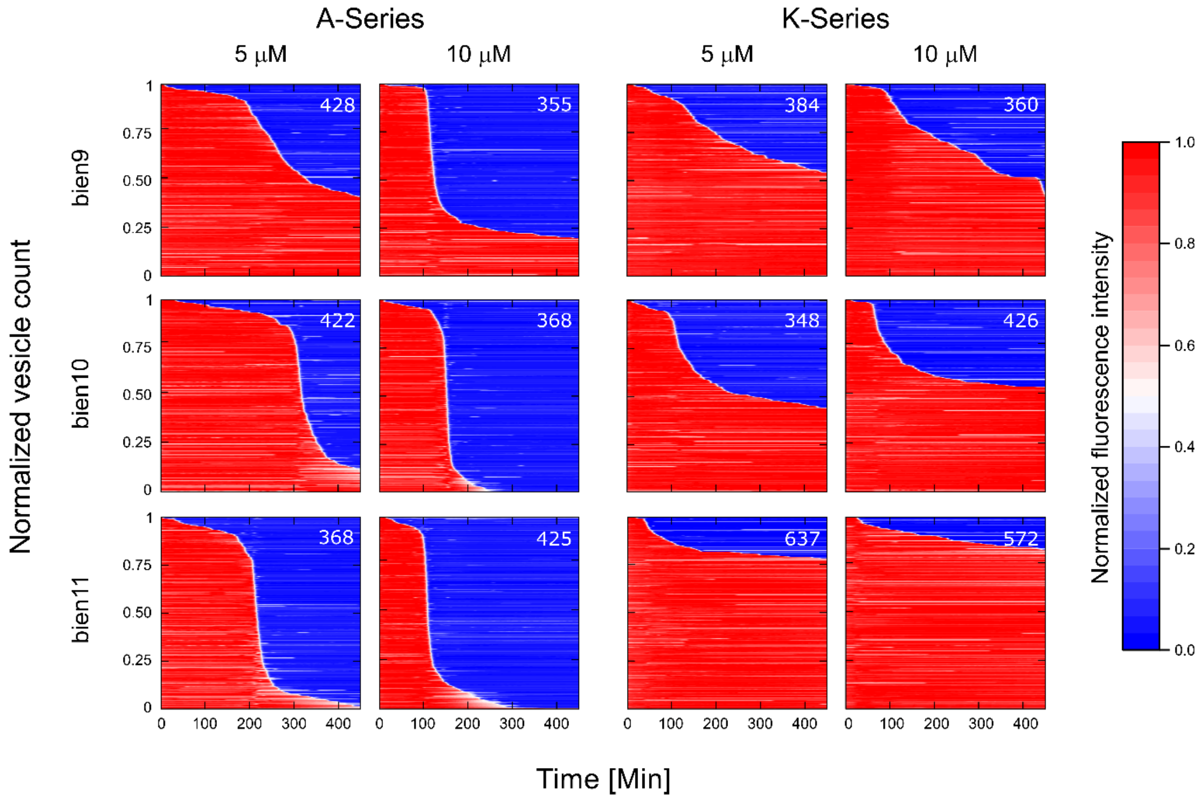

**Figure S3. A summary of the membranolytic activity of the bien peptides on populations of bacterial membrane-mimicking lipid vesicles. Dataset 2 of 2.** For completeness, includes data that has been reprinted (adapted) with permission from K. Hammond, F. Cipcigan, K. Al Nahas *et al.*, "Switching Cytolytic Nanopores into Antimicrobial Fractal Ruptures by a Single Side Chain Mutation", *ACS Nano*, 15 (6), 9679-9689 (2021). Copyright 2021 American Chemical Society. Trapped vesicles were continuously treated with the peptides at 5 μM and 10 μM concentrations and their morphology was observed overnight. Each horizontal line depicts the locally normalized intensity of the fluorescent dye HPTS encapsulated in a single trapped vesicle (with global background subtraction) over time. The vesicle's membrane is considered intact at high fluorescence intensity (red) and compromised at low fluorescent signal (blue). The intensity traces were ordered by the critical viability time point, which is defined as the point when the fluorescence intensity of a vesicle decreases below 50% of its initial intensity. The total number of analysed vesicles is reported in white in the top right corner of every plot. The results show that the bienA series of peptides is membranolytic, with potency increasing as one progresses from bienA9 to bienA11 and with an increase in the respective drug concentrations. However, the bienK series is not obviously membranolytic – there appears to be a weakening of the membranes in relation to controls, but there is no obvious concentration dependence, and the trend is reversed with respect to the bienA series, with bienK11 being the least potent.

**Supplementary references:**

(1) Goode, O.; Smith, A.; Zarkan, A.; Cama, J.; Invergo, B. M.; Belgami, D.; Caño-Muñiz, S.; Metz, J.; O'Neill, P.; Jeffries, A.; Norville, I. H.; David, J.; Summers, D.; Pagliara, S. Persister *Escherichia coli* Cells Have a Lower Intracellular pH than Susceptible Cells but Maintain Their pH in Response to Antibiotic Treatment. *mBio* **2021**, *12* (4), e00909-21.

(2) Goode, O.; Smith, A.; Łapińska, U.; Bamford, R.; Kahveci, Z.; Glover, G.; Attrill, E.; Carr, A.; Metz, J.; Pagliara, S. Heterologous Protein Expression Favors the Formation of Protein Aggregates in Persister and Viable but Nonculturable Bacteria´. *ACS Infect. Dis.* **2021**, *7*, 1848–1858.

(3) Bamford, R. A.; Smith, A.; Metz, J.; Glover, G.; Titball, R. W.; Pagliara, S. Investigating the Physiology of Viable but Non-Culturable Bacteria by Microfluidics and Time-Lapse Microscopy. *BMC Biol.* **2017**, *15*, 121.
